# Dynamic shape remodeling of vesicles by internal active filaments

**DOI:** 10.1101/2025.07.16.665046

**Authors:** Arash Karaei Shiraz, Amir H. Bahrami

## Abstract

To interact with their environment, living cells use active cytoskeletal forces to form dynamic membrane structures such as tubular filopodia and sheet-like lamellipodia. To understand the formation and dynamics of these structures, we perform non-equilibrium simulations of dynamically triangulated vesicles under osmotic stress. We investigate vesicle shape remodeling driven by local effects of internal active filaments, as well as large-scale shape transformations resulting from volume changes controlled by osmotic pressure. We identify the morphological behavior of vesicles across varying volumes and filament properties, including concentration, mobility, stiffness, and length. Our simulations reveal dynamic, unstable vesicle structures—such as branched tubes, sheet-tubes, cup-tubes, and compartmentalized vesicles—composed of tubular, sheet-like, and cup-like segments. These structures continuously reorganize, interconverting between different shape components while maintaining nearly constant proportions. In particular, unstable branched tubes form under low vesicle volume and low filament mobility. Remarkably, their restructuring accelerates as filament mobility decreases, suggesting that their dynamics are primarily governed by global vesicle remodeling under osmotic stress. Notably, branched tubes arise only in the presence of active filaments and vanish when filaments become apolar due to shortening and loss of anisotropy. Our findings reveal novel non-equilibrium pathways for generating unstable, dynamic cellular structures such as branched tubes, sheet-tubes, and compartmentalized vesicles. These insights not only advance our understanding of complex organelle morphologies and cellular protrusions but also suggest new mechanisms for actively shaping synthetic membrane systems.

---

Active matter systems investigate how the collective motion of self-propelled microscale elements gives rise to macroscopic emergent phenomena, with broad applications in natural, biological, and materials sciences.^1–5^ Among these, biological active matter is fundamental to cell biology, where energy-consuming elements drive cellular organization and function.^4,6^ Moreover, controlling active forces enables the development of synthetic materials with life-like properties, not only for engineering applications such as soft robotics^7–9^ but also for designing biomimetic systems that mimic living organisms, with broad applications in synthetic biology.^10,11^

Biological active matter focuses on living organisms, which rely on the hierarchical selforganization of energy-consuming active components that operate away from equilibrium to fulfill their functions.^4,6,12^ In particular, living cells regulate their structure by harnessing internal active forces from the cytoskeletal network,^13,14^ which drive deformations in lipid membrane boundaries. The resulting active membrane remodeling is essential for shaping various non-equilibrium cellular structures.^15,16^ Prime examples include highly branched dendritic networks in neurons,^17,18^ membrane protrusions such as sheet-like lamellipodia and finger-like filopodia in cell migration,^15,19–21^ cytoplasmic streaming,^22^ and active processes like endocytosis and phagocytosis.^23,24^ Other examples include cellular blebs and tethers,^25,26^ as well as dumbbell-like structures that emerge during cytokinesis and cell division.^27–29^ Membrane deformation by active forces is central to all these phenomena.

Studying active membrane dynamics aims to elucidate how interactions with the active cytoskeletal network shape cellular membranes. However, theoretical and computational studies of these interactions remain highly complex. A bottom-up approach in synthetic biology^30^ addresses this challenge by using lipid vesicles ^16,25,31,32^ and liquid droplets^33,34^ as biomimetic models to investigate cellular processes with controlled complexity.^4^ Active vesicles, in particular, provide minimalistic cell models for studying these processes experimentally and computationally.

Non-equilibrium active vesicles can exhibit shapes that resemble well-known stable configurations—prolate tubes, oblate sheets, and stomatocytes (cups)^35–37^—or adopt intrinsically unstable structures such as branched tubes, sheet-tubes (tubes invaginated from sheets), cup-tubes, and pearled tubes, which are abundant in living cells. ^38,39^ These unstable structures require the presence of active agents or curved proteins to maintain their shape. Active forces not only sustain these unstable shapes but also stabilize sheets, tubes, and cups under conditions where they would otherwise be unstable.^40,41^ For instance, they can maintain otherwise unstable membrane tubes at low volume-to-area ratios, ^42^ where cups represent the stable shapes.^35–37^

Depending on their driving mechanism, non-equilibrium shape transformations of active vesicles arise from the interplay between global, large-scale deformations and local, shortrange remodeling. Local factors, such as the swim pressure of active particles, directly induce localized shape changes in vesicle membranes.^43–45^ In contrast, global factors such as osmotic pressure and membrane area expansion through membrane growth^46^ indirectly sculpt vesicles by regulating their volume-to-area ratio. Similar global boundary effects have been observed in the bulk behavior of active systems. ^47^ Moreover, membrane-mediated alignment of anisotropic active elements has been shown to drive system-spanning global shape transformations.^42,48^ More recently, the interplay between the active cytoskeletal network and the deformable membrane has been shown to produce large-scale, traveling membrane deformations in synthetic cells.^49^

Active shape remodeling of vesicles with internal particles depends on particle properties, particle-membrane interactions, and osmotic pressure, which regulates the vesicle volume-to-area ratio. Membrane properties, such as elasticity and spontaneous curvature, play a crucial role. Particle properties include the mobility or propulsion strength of the self-propelled active elements, and in the case of filaments, their anisotropy and bending deformability.^48,50,51^ Theoretical analyses have explored the dynamic shape fluctuations of active membranes.^52,53^ Shape transformation of vesicles induced by isotropic active particles—both with^50^ and without membrane adhesion^16,43,50,52,54^—as well as by anisotropic active filaments,^10,48,55^ are relatively well understood. Similar studies have focused on 2D simulations of active vesicles.^10,56–58^ Nevertheless, dynamic shape transformations of active vesicles are poorly understood. Recent studies are limited to pearled tubes, which dynamically vary their compartment number due to the swim pressure of internal active beads.^43^ However, the general dynamic shape remodeling of active vesicles—particularly how largescale vesicle transformations driven by osmotic pressure, combined with the local effects of anisotropic active filaments, sculpt vesicles at low volume-to-area ratios—remains largely unexplored.

Here, we explore non-equilibrium dynamic shape transformations of elastic vesicles driven by osmotic pressure and internal self-propelled, semi-flexible active filaments. Using over-damped Langevin dynamics simulations of dynamically triangulated vesicles, we map the morphological behavior of active vesicles as a function of filament properties—including mobility, concentration, length, and flexibility—as well as the vesicle volume-to-area ratio controlled by osmotic stress. We observe dynamic sheet-like, tubular, and cup-shaped vesicles, along with unstable hybrid structures such as sheet-tubes, branched tubes, and pearled tubes. Unlike previous studies where vesicles with internal active filaments reached steadystate morphologies in the absence of osmotic stress,^48^ we observe vesicles that dynamically reorganize their structures, highlighting the critical role of osmotic pressure in large-scale vesicle remodeling. Remarkably, at low volume-to-area ratios, vesicles form highly branched tubular networks, with tube dynamically emerging and retracting at rates proportional to filament mobility and concentration. Interestingly, lower filament concentrations promote more dynamic reorganization. Finally, we demonstrate that filament length and flexibility, which together define filament anisotropy, are key determinants of vesicle morphology.

## Results and Discussion

The morphological behavior of non-equilibrium vesicles under osmotic stress and internal active filaments is governed by a combination of filament and vesicle variables, including the vesicle volume-to-area ratio, filament mobility (activity), concentration, and anisotropy, as determined by filament length and bending stiffness. Accordingly, vesicle shapes are controlled by dimensionless parameters: the reduced volume 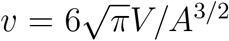 of the vesicle, where *A* and *V* are the vesicle surface area and volume, respectively; the Péclet number Pe = *σf*_*p*_*/k*_*B*_𝕋 ∈ [3.75, 15]; the filament concentration *ϕ*; the aspect ratio (rescaled length) ℒ of the filaments; and the rescaled filament bending stiffness *χ* = *κ*_*f*_ */κ*. Here, *f*_*p*_ is the propulsion force, *k*_*B*_ is the Boltzmann constant, and 𝕋 is the temperature. In our model, active filaments are represented as polymer chains composed of isotropic spherical beads of size *σ*, sequentially linked by harmonic bond potentials. For a given number of filaments (*N*_*f*_), the filament concentration, *ϕ*, is defined as the volume fraction of a spherical vesicle (*v* = 1) that is occupied by all filament beads. All simulations are performed in the constant-area ensemble, with the vesicle area *A*_0_ = 4*πR*^2^ defining the size scale *R*. The membrane has a bending rigidity *κ*, and *κ*_*f*_ denotes the bending stiffness of the filaments, see Methods for more details.

We find that non-equilibrium vesicles exhibit complex dynamic morphologies, adopting either one of the stable vesicle shapes—tubes (T), sheets (S), or cups (C), also observed for equilibrium vesicles^35,36^—or hybrid forms that combine these structures, Fig. 1A. The unstable hybrid shapes include sheet-tubes (ST), which combine tubular and sheet-like sections with tubes protruding from the sheets; branched tubes (BT), featuring three-way tubular junctions; and compartmentalized vesicles (CV), which comprise either pear-shaped vesicles that combine spherical and tubular sections or compartmentalized structures with partial constrictions forming interconnected, dynamic compartments, Figs. 1A and S1. Membrane asymmetry, Δ*a*, represents the area difference between the two membrane leaflets. For a given *v*, Δ*a* assumes distinct values for tubes (Δ*a*_*T*_), sheets (Δ*a*_*S*_), and cups (Δ*a*_*C*_).^35,37^ For combined vesicle shapes, Δ*a* continuously varies between these characteristic values. For example, varying Δ*a* between Δ*a*_*S*_ and Δ*a*_*T*_ results in sheet-tubes with different proportions of tubular and sheet-like sections. Therefore, we use the dimensionless shape index, *α* = Δ*a/*Δ*a*_*T*_, to characterize vesicle morphology, where *α* = 1 corresponds to tubular structures for a given *v*, see Methods.

**Figure 1.**
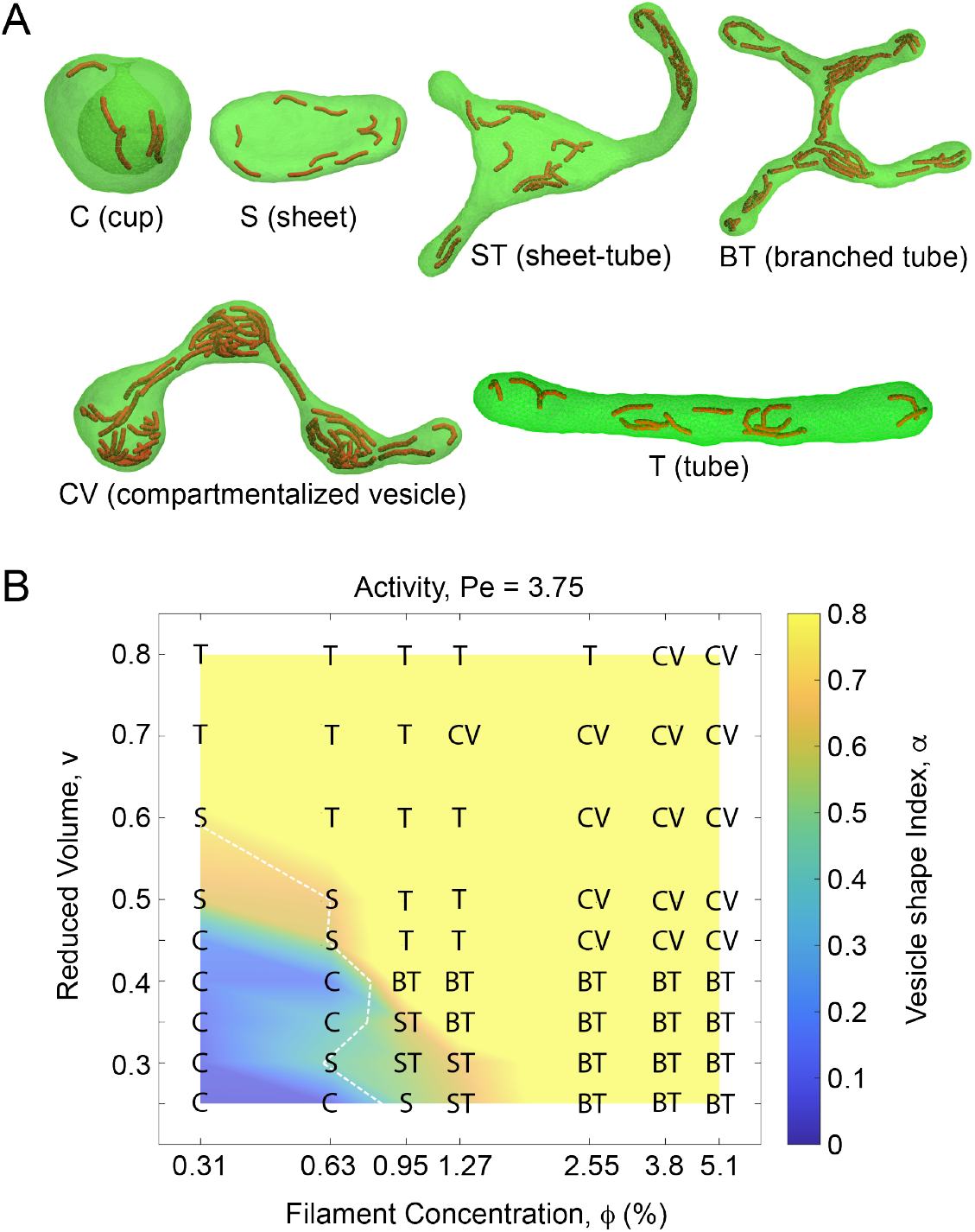
Vesicle morphologies formed by internal active filaments. (A) Active vesicles either adopt one of the fundamental vesicle shapes—tubes (T), sheets (S), or cups (C)—which are also observed at equilibrium, or form hybrid structures such as sheet-tubes (ST), branched tubes (BT), and compartmentalized vesicles (CV) including pear-shaped vesicles and pearled tubes. (B) Morphology diagram of active vesicles with varying *ϕ* and *v* at a fixed activity Pe = 3.75 for relatively stiff filaments (*χ* = 25) of length (ℒ = 6.12). Vesicle shapes are characterized by the shape index *α*, which is based on membrane asymmetry (Δ*a*) and is color-mapped onto the shape diagram. The white dashed line indicates toroidal sheets predicted by the theoretical vesicle model at different values of *v* (see Methods).

For a given Pe and fixed filament properties, χ and ℒ, the morphological behavior of active vesicles depends on *v* and *ϕ*, as shown in Fig. 1B. The morphology diagram can be divided into four distinct regimes based on vesicle structures. At low values of *v* and *ϕ*, vesicles adopt cup shapes, represented by the lower-left (blue) region of the diagram. Slight increases in *ϕ* and *v* give rise to two distinct regimes within the green region. At low *ϕ*, increasing *v* promotes the formation of sheets (S) in the upper-left part of the green region (brown), which rapidly transition into tubes beyond the white dashed lines representing theoretically predicted toroidal sheets. Further increasing *v* establishes the tubular regime in the top-left part of the diagram, characterized by simple tubes (T) with nearly uniform cross-sections.

In contrast, increasing *ϕ* while keeping *v* low drives a rapid transition from cups to sheet-tubes (ST), located in the lower-right brown region of the green area. Interestingly, further increasing *ϕ* transforms these sheet-tubes into branched tubes (BT) with a dynamically varying number of tubular junctions, a regime uniquely observed at low *v* and high *ϕ* in the bottom-right corner of the diagram. Finally, a simultaneous increase in *v* and *ϕ* leads to the fourth regime, located at the top-right of the diagram, predominantly consisting of dynamically reorganizing compartmentalized vesicles (CV) with constricted membrane necks separating distinct compartments. White dashed lines in Fig. 1B represent theoretical toroidal sheets at each volume. These lines closely follow the simulated sheet-like vesicles, indicating that *α* serves as a useful measure for distinguishing vesicle shapes. Interestingly, branched tubular networks are observed only at sufficiently small values of *v* ≤ 0.4 and relatively large *ϕ* ≥ 0.95, while larger values of *v* typically give rise to tubes and compartmentalized vesicles. Consistent with previous studies of active vesicle remodeling in the absence of osmotic effects, ^48^ we find that filament mobility does not significantly influence the morphological behavior of active vesicle. While the four vesicle regimes remain similar with slightly shifted boundaries, increasing Pe from 3.75 to 15 decreases the abundance of branched tubes, converting them into tubular vesicles, Fig. S2.

The active vesicles we observe here dynamically reorganize their structures. In the absence of osmotic stress, active vesicles with confined filaments have been shown to exhibit steady-state morphologies.^48^ In contrast, we observe that under osmotic stress, they undergo dynamic shape transformations, particularly evident in unstable hybrid structures such as sheet-tubes, branched tubes, and compartmentalized vesicles. For instance, individual tubes within branched tubular networks emerge and retract dynamically, resulting in a variable number of three-way tubular junctions. This complex dynamic behavior depends on system parameters and is consistently observed across different values of *v* in the branched tubular regime, Figs. 2A,B and S3A-E, and Movies S1 and S2. Similarly, the sheet-like and tubular sections of sheet-tubes dynamically interconvert, while maintaining a relatively constant proportion of each, Figs. 2D and S3I, and Movie S3. Consequently, the total length of the tubular segments, as well as the tube diameter, remains nearly constant despite fluctuations in the number and individual lengths of the tubes. Comparable dynamic reorganization is observed in compartmentalized vesicles, where both the shape and number of internal compartments fluctuate over time, as well as in pear vesicles with relatively constant proportions of spherical and tubular sections, Figs. 2E and S3G. Even fundamental shapes such as tubes, sheets, and cups exhibit continuous remodeling while preserving their overall geometry, as demonstrated by the temporal evolution of active sheets, Figs. 2F.

**Figure 2.**
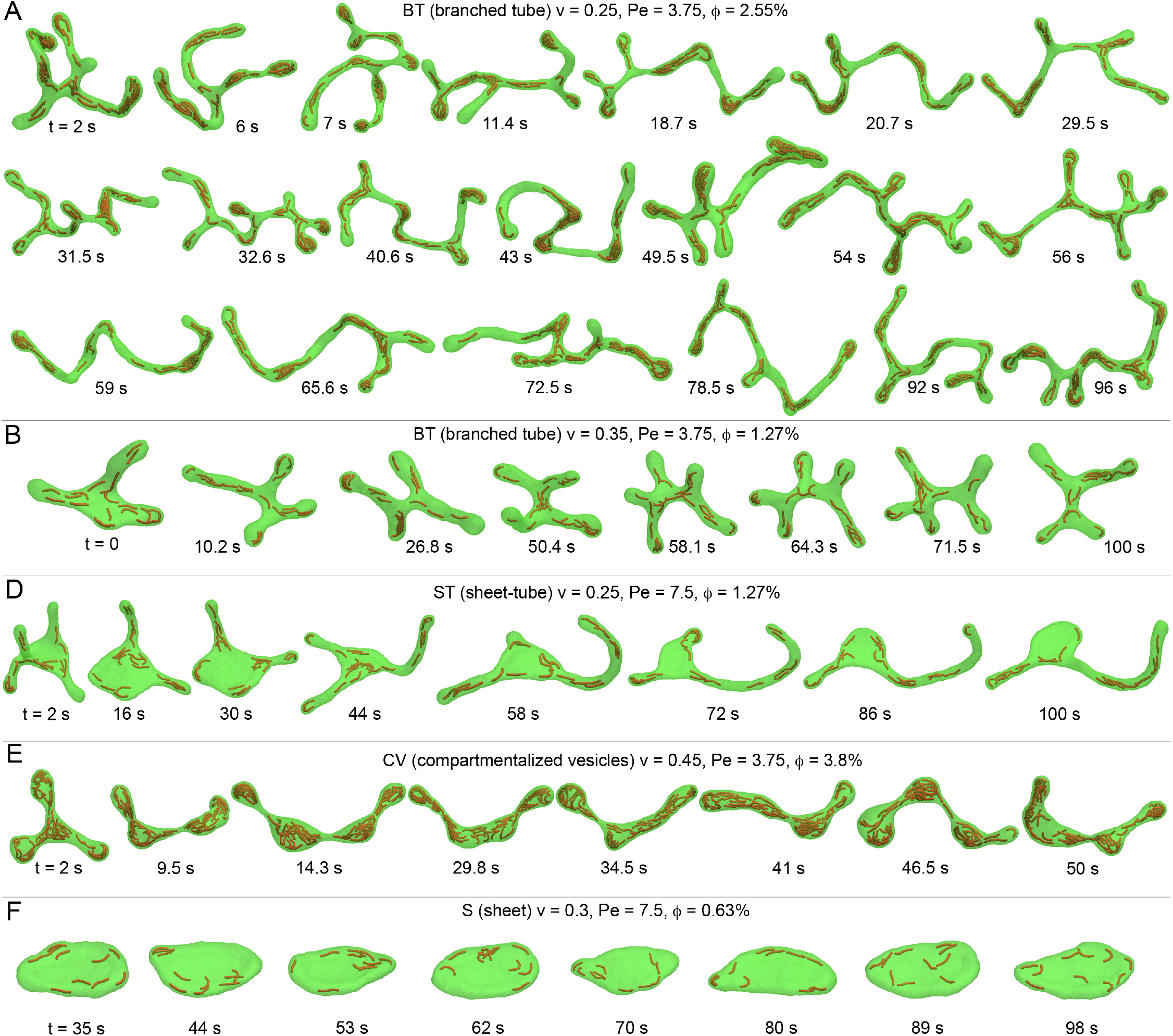
Temporal evolution of dynamically reorganizing active vesicles. Branched tubes with relatively small volumes, (A) *v* = 0.25 and (B) *v* = 0.35, continuously reorganize their structures by extending and retracting tubes, forming a variable number of three-way tubular junctions. Movies S1 and S2 show real-time animations of panels A and B, respectively. (D) A dynamic hybrid sheet-tube composed of interconnected sheet-like and tubular sections that interconvert while maintaining a nearly constant fraction of each section over 100 seconds of simulation. The length of the tubular sections and the area of the sheet-like section remain almost unchanged. Movie S3 shows a real-time animation of the sheet-tube structure in panel D. (E) Dynamic evolution of a compartmentalized vesicle, composed of restructuring compartments, at a relatively large *φ* over 50 seconds. (F) A sheet with relatively low *φ* and *v*, which retains its sheet-like morphology throughout the 100-second simulation, despite continuous structural reorganization.

To better understand the dynamic reorganization of active vesicles and its relation to *v* and *ϕ*, we focus on the branched tubular networks, which are particularly prominent at low *v*, Fig. 2A,B. The fundamental building block of these networks is a three-way tubular junction, where two tubes intersect, referred to hereafter as a tubular junction. The top-right panel of Fig. 2A shows two such junctions connected to each other to form an H-shaped configuration. Each junction consists of two intersecting tubes with nearly identical diameters. In living cells, similar unstable tubular junctions can emerge through the equilibrium fusion of two membrane tubes.^59^ In our simulations, they form via the non-equilibrium sprouting of tubular protrusions driven by active forces from self-propelled filaments. Conversely, a tubular junction vanishes when one of the two intersecting tubes retracts. The number of tubular junctions determines the network complexity, while its variation over time provides a measure for the network dynamics.

We monitored the temporal evolution of the number of tubular junctions, Figs. 3A and S4 and Movies S1 and S2. We then calculated the rate of change in network junctions, *r* = *r*_*e*_ + *r*_*r*_, as the sum of the emergence (*r*_*e*_) and retraction (*r*_*r*_) rates of tubular junctions per second for branched tubes with varying *v* and *ϕ*, Fig. 3A. For sufficiently low *v* = 0.25, we observed branched tubular networks composed of 1 to 5 tubular junctions. At a constant Pe, we found that *r* almost always increases with *ϕ*, Fig. 3B. Interestingly, however, *r* decreases as Pe increases at a fixed *ϕ* (Fig. 3D) implying that tubular network reorganization occurs more frequently for filaments with lower mobility. This trend persists across different values of *ϕ* ≥ 2.55 in the branched tubular regime.

**Figure 3.**
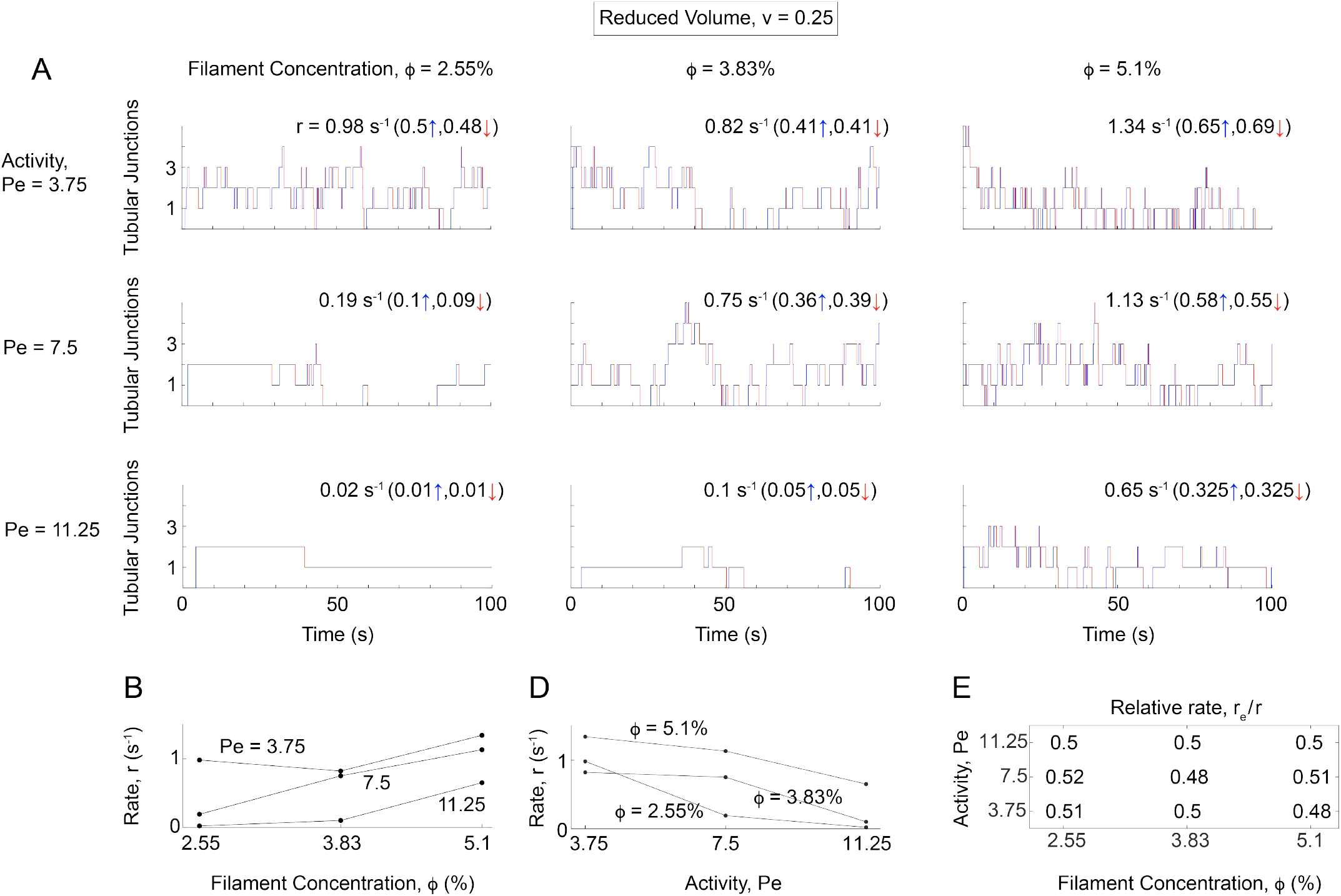
Dynamic reorganization of branched tubes. (A) Temporal evolution of the number of three-way tubular junctions for different values of Pe and *ϕ* at *v* = 0.25. The total rate of change in the number of tubular junctions is given by the sum of the tube emergence rate (blue lines) and the tube retraction rate (red lines). (B) The rate of change in tubular junctions increases with *ϕ* for a fixed Pe, while (D) it decreases with Pe at a constant *ϕ*. (E) All branched tubes exhibit nearly identical emergence and retraction rates, *r*_*e*_ ≈ 0.5, implying that tube emergence and retraction occur with almost equal frequency.

The tubes emerge and retract with almost identical rates for all values of Pe and *ϕ*, as observed in the ratio *r*_*e*_*/r* ≈ 0.5 for all networks, Fig. 3E. Therefore, despite the absence of steady-state behavior, non-equilibrium tubular networks maintain their complexity while dynamically reorganizing their structure. This novel feature arises from the combined effects of local vesicle remodeling driven by filament mobility and global large-scale shape transformations induced by osmotic stress. A comparison between Figs. 3A and S4 highlights the significant influence of *v* on the dynamics of tubular networks. A slight increase in *v* from 0.25 to 0.3 leads to markedly slower reorganization of branched tubes, as reflected in lower values of *r*.

In addition to osmotic stress and filament mobility, vesicle remodeling is strongly influenced by the mechanical properties of the filaments, particularly their length and stiffness, ℒ and χ. The rescaled filament length, ℒ, corresponds to the equilibrium filament length and is thus proportional to *N*_*bf*_ − 1, where *N*_*bf*_ is the fixed number of filament beads in each simulation. For filaments interacting with vesicles, the actual filament length depends on χ and ℒ, as well as *ϕ* and *v*. Consequently, the average end-to-end distance—commonly referred to as the filament persistence length, P—provides a more accurate measure for distinguishing anisotropic (polar) filaments from isotropic (apolar) beads.

At *v* = 0.25, we varied χ within the interval [0, 25], with *χ* directly controlling the filament anisotropy via changing P. As expected, vesicle structures strongly depend on χ, and thus on filament anisotropy, as shown in Fig. 4A for Pe = 3.75. For relatively stiff filaments with high χ = 25, we recover our previous results shown in Fig. 1B. However, decreasing χ, which corresponds to more flexible and less straight filaments, drastically changes the morphological behavior of the vesicles. As χ decreases, the transition from cups to sheets and sheet-tubes and eventually to branched tubes, occurs at progressively higher values of *ϕ*. Consequently, branched tubes, predominantly observed at high *ϕ* and χ, gradually diminish as χ decreases from 25 and eventually vanish at χ = 10. For sufficiently low χ *<* 10, where filaments lose their anisotropy and crumple into apolar lumps, only cup-shaped vesicles are observed, even at high *ϕ*, Fig. 4A. A similar behavior is observed for higher Pe values (Pe ≥ 7.5), Fig. S5. Consistently, corresponding values of P show apolar crumpled filaments for χ*<* 10, Fig. 4B.

**Figure 4.**
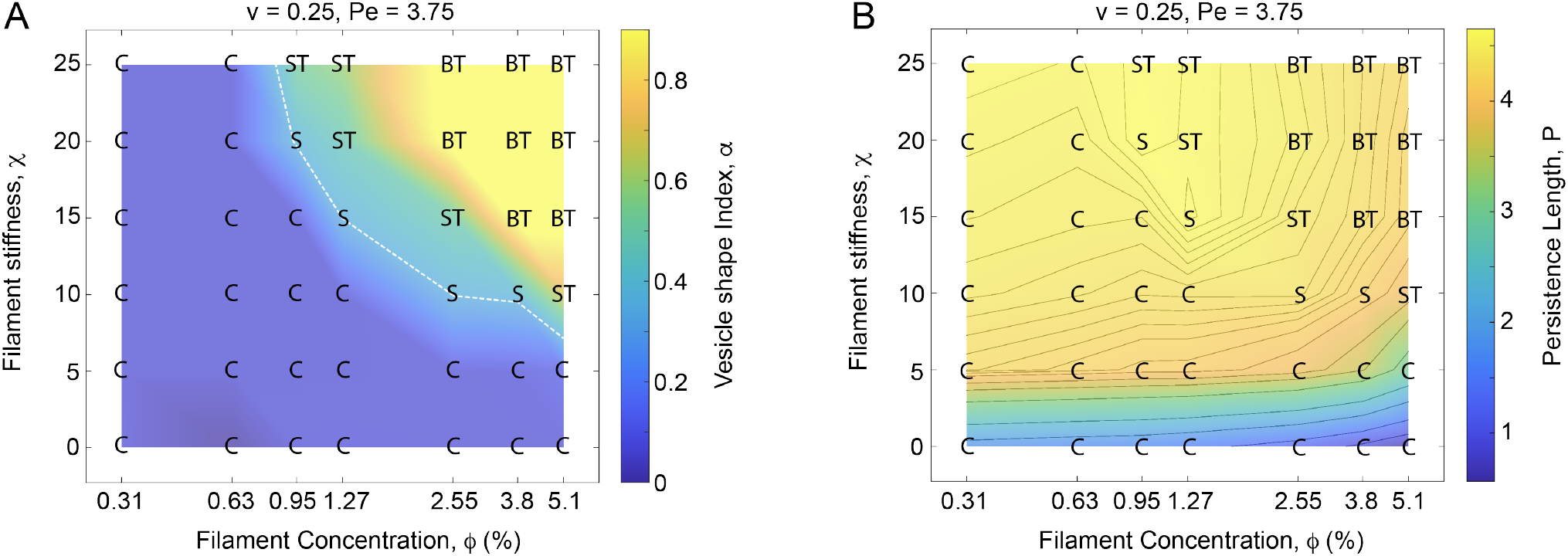
Vesicle morphologies under filament bending stiffness. (A) Morphological behavior of vesicles with active filaments at varying filament stiffness. Lower filament stiffness delays the transition from cups to sheets, sheet-tube structures, and eventually branched tubes at larger values of *ϕ*. Reduced stiffness also leads to the formation of crumpled filament aggregates, transforming branched tubes into cups for χ ≤ 5. The white dashed line indicates toroidal sheets predicted by the theoretical vesicle model (see Methods). (B) For small *χ*, crumpled filaments exhibit low persistence lengths, P, in cups, consistent with the vesicle shapes shown in panel A.

For fixed *v* = 0.25 and χ = 25, we then varied *N*_*bf*_ within [1, 10] to form filaments with 1 ≤ ℒ ≤ 12.52 at different values of Pe = 3.75, 7.5, 11.25 (Fig. 5). The lower limit ℒ = 1 with *N*_*bf*_ = 1, corresponds to active vesicles containing internal apolar spherical beads. The morphology diagram of active vesicles for varying *ϕ* and ℒ features an intermediate branched tubular regime (yellow) surrounded by two non-tubular regimes from below and above. Two transition regions (orange bands) separate the non-tubular regimes from the branched tubular regime. At low ℒ ≲ 4, below the nearly horizontal transition band (orange), only cups are observed across different values of *ϕ* and Pe. As ℒ increases, sheets and sheet-tubes emerge along and near the transition band, rapidly evolving into branched tubes beyond it. Interestingly, further increasing ℒ results in the recovery of sheets and sheet-tubes and eventually cups at sufficiently high Pe beyond a second transition band (oragne) as seen near the top left of the morphology diagrams. The lower transition band is closely approximated by the lower white dashed lines, which correspond to toroidal sheets. For sufficiently high Pe = 11.25, the upper transition band also encompasses the top white dashed line corresponding to toroidal sheets in the right panel of Fig. 5.

**Figure 5.**
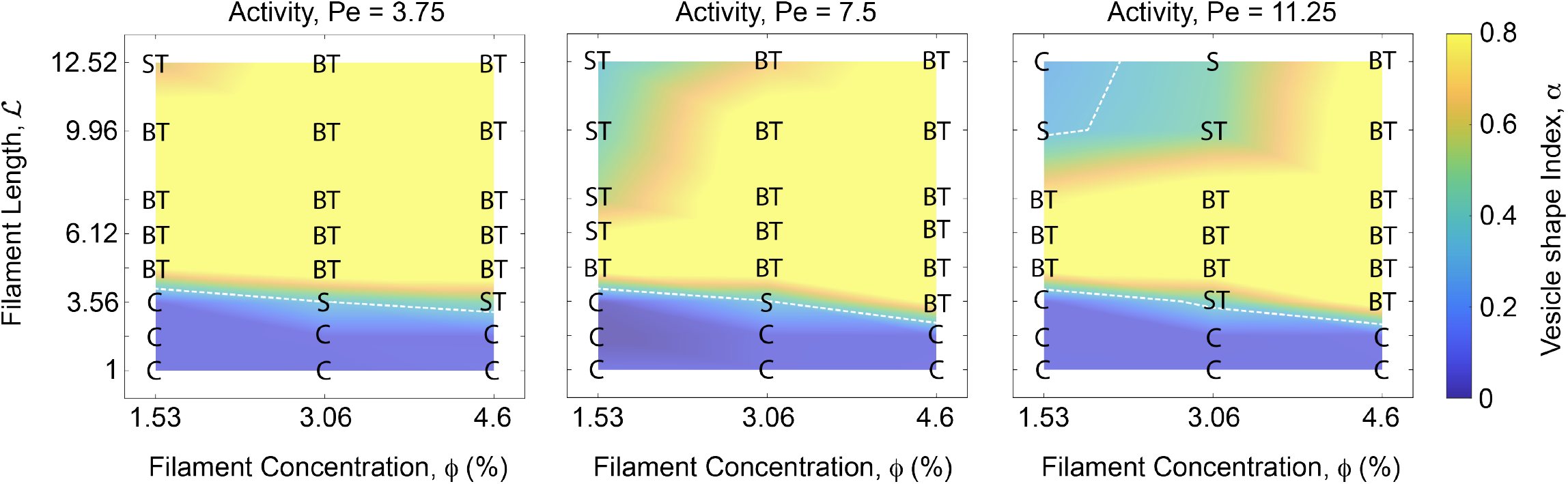
Vesicle morphologies under filament length. The morphological behavior of active vesicles exhibits a Pe-dependent response with varying filament length, ℒ. Branched tubular structures (yellow region) appear only within a specific intermediate range of ℒ, beyond which they transition into either cups or sheets. The white dashed lines indicate toroidal sheets predicted by the theoretical vesicle model at different values of *v* (see Methods). The intermediate branched tubular regime diminishes with increasing Pe. The simulations are performed at *v* = 0.25 and χ = 25.

The transitions between the branched tubular regime and the surrounding non-tubular regimes exhibit different behaviors depending on Pe. While the bottom transition bands remain nearly horizontal and largely independent of *ϕ*, the top transition band shows a strong dependence on *ϕ*. Consequently, the initially small top non-tubular regime, confined to low *ϕ* ≲ 2.5 at Pe = 3.75, expands with increasing Pe, reaching *ϕ* ≲ 4 at Pe = 11.25. As a result, the top transition band, which is absent at Pe = 3.75 and 7.5 due to the lack of sheets, emerges at Pe = 11.25 (right panel of Fig. 5). Therefore, vesicle shapes exhibit non-monotonic behavior with respect to ℒ, making their morphological transitions highly sensitive to Pe. This distinct sensitivity to Pe is unique to variations in ℒ, in contrast to *χ* and *v*, where vesicle shapes show only a weak dependence on Pe. Our results indicate that tubular membranes, particularly dynamically reorganizing branched tubes (BT), form within an intermediate range of ℒ, which narrows as Pe increases. For very short filaments below this range and very long ones beyond it, branched tubes are not observed.

In addition to different dynamic vesicle structures, we rarely observe short-lived membrane tethers with life times ≲ 1s, which occur almost exclusively for branched tubular networks with relatively large *ϕ*, Fig. 6. These long, narrow tethers have diameters comparable to the filament width (i.e., the size of a single bead) and are typically thinner than the parent tube from which they extend. They usually broaden and reintegrate into the original tube within a short time. Similar tethers have been reported in vesicles with both normal and sticky active beads.^16,50,51^

**Figure 6.**
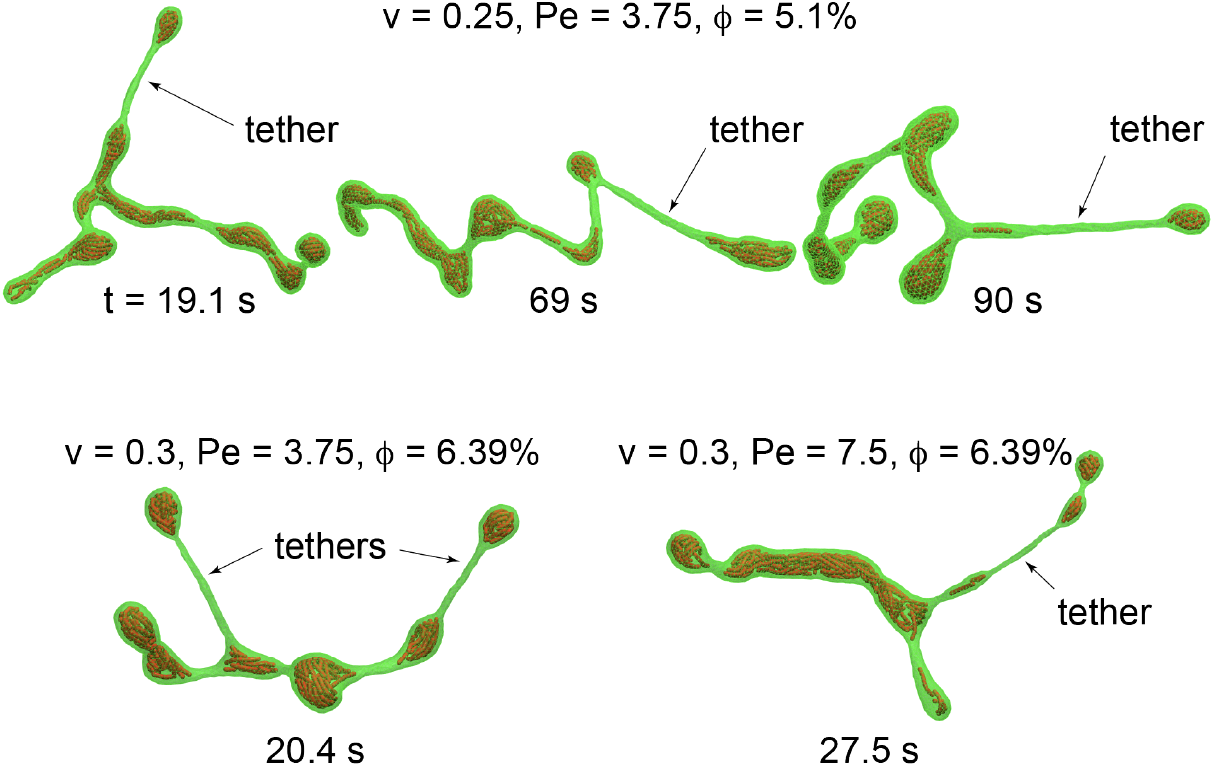
Membrane tethers form at high filament concentrations, *ϕ*. Short-lived tethers (lifetime ≲ 1 s) are observed almost exclusively on branched tubes with relatively large *ϕ*. These tethers have smaller diameters than their parent tubes.

For less anisotropic filaments with relatively low P—corresponding to small ℒ or χ—we also observe additional dynamically unstable cup-like variants. For instance, increasing *ϕ* at *v* = 0.25 leads to the formation of multi-compartmentalized cups, Figs. S6A and S6B (top-left panel). Another example is tube-cups, which consist of both cup-like and tubular sections and are observed at low ℒ, Fig. S6B, bottom panel. Like other unstable hybrid structures, such as sheet-tubes, branched tubes, and compartmentalized vesicles, these shapes also undergo continuous morphological reorganization, with shape units dynamically emerging, disappearing, and interconverting.

To explore how vesicle shapes and their dynamics relate to the large-scale effects of external osmotic stress and the local effects of internal active filaments, we calculate the average membrane tension 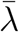 in active vesicles (see Methods) and compare it to *λ*_0_ = *R*^2^*k*_*B*_𝕋*/*(*πσ*^4^). The vesicles simulated here exhibit membrane tension in the range 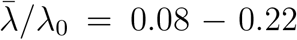, corresponding to approximately 1 − 3 *μN/m* for a typical vesicle size of *R* = 1 *μm*. These vesicles are thus classified as floppy vesicles with almost negligible membrane tensions. While 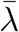 appears to increase with *ϕ*, it remains constant across different values of Pe ∈ [3.75, 15] (Fig. S8), suggesting that the shapes and dynamics of these floppy vesicles in the low-tension regime are not primarily governed by membrane tension.

## Methods

Our model is composed of dynamically triangulated vesicles and active filaments that move inside the vesicles under external active forces.

### Active filaments

Active filaments are polymer chains composed of multiple active beads, modeled as active Brownian particles. Each two beads belonging to different filaments experience a repulsive force with the interaction potential given by

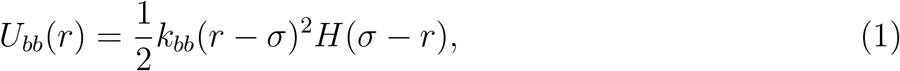

where *r* is he distance between the beads, *k*_*bb*_ is the potential coefficient adjusting the repulsion strength, *σ* denotes the effective bead diameter, and *H* is the heaviside step function.^43^ Each filament consists *N*_*bf*_ beads which are connected by *N*_*bf*_ − 1 links through applying a harmonic bond potential:

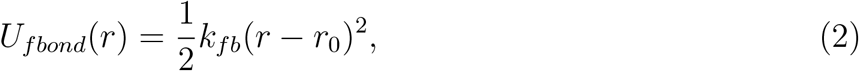

between every two consecutive beads inside the filament. The bond stiffness, *k*_*fb*_, and the equilibrium bond length *r*_0_ control the bond lengths of the filaments. The filament length is defined as ℒ = (*r*_0_(*N*_*bf*_ − 1) + *σ*)*/σ*. Bending deformations of the filaments is governed by a bending potential given as:

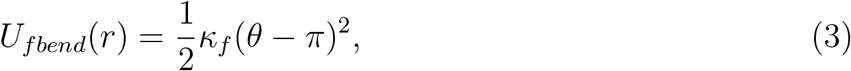

with *k*_*f*_ representing the filament bending stiffness and *θ* denoting all the *N*_*bf*_ − 2 angles formed between each three consecutive beads in each filament. The persistence length P characterizes the effective end-to-end filament distance, rescaled by *σ*. A total number of *N*_*f*_ filaments in vesicles results in a filament concentration, *ϕ* = *N*_*f*_ *N*_*bf*_ (*σ/*2*R*)^3^, representing the volume fraction of a spherical vesicle with surface area *A* that is occupied by all filaments. Interactions between the filaments and the vesicle are governed through repulsive forces acting between all filament beads and the vesicle nodes (vertices) given by

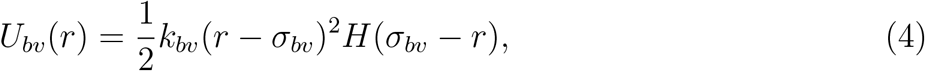

where *r* is the bead-vertex distance, *k*_*bv*_ = *k*_*bb*_ is the potential coefficient adjusting the repulsion strength, and *σ*_*bv*_ = *σ* denotes the effective bead-vesicle distance.

Filament displacement is governed by the motion of filament beads with a position vector **r**_*i*_(*i* = 1..*N*_*bf*_) following the Langevin equation,

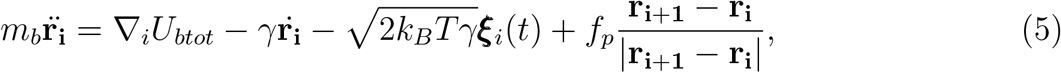

where *U*_*btot*_ is the sum of all interaction potentials acting on beads, *m*_*b*_ is the bead mass, 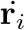 and 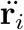 are the first and the second time derivatives of the position vector **r**_*i*_ of bead *i*, ∇ is the spatial derivative at the position vector of bead *i*. Here, *γ* represents the effect of the surrounding viscous fluid on the filaments and ***ξ***_*i*_(*t*) is a Gaussian white noise with its Cartesian components following ⟨*ξ*_*i*_(*t*) ⟩ = 0 and ⟨*ξ*_*i*_(*t*)*ξ*_*j*_(*t*^′^) ⟩ = *δ*_*ij*_*δ*(*t* − *t*^′^) representing thermal fluctuations. The last term in Eq. 5 represents the external active force of magnitude *f*_*p*_, applied to filament beads along the filament.

Langevin simulations were performed for 50 seconds and, in some cases, extended to 100 seconds. Data was collected every 0.01 seconds, and the last 30 seconds of each simulation were used to calculate the mean and standard deviation of various properties.

### Triangulated membrane model of the vesicles

The vesicles in our model are presented by a three dimensional triangulated membrane composed of *N*_*v*_ point vertices or nodes connected by *N*_*l*_ links which together form *N*_*t*_ = 2(*N*_*v*_ − 2) triangles.^60^ Membrane bending is governed by the Helfrich model of curvature elasticity^61^ according to which the bending energy of the vesicle, with negligible spontaneous curvature, *C*_0_ = 0, is given by:

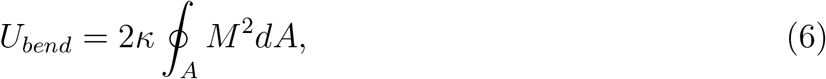

where *M* = (*C*_1_ + *C*_2_)*/*2 is the mean of the two principle curvatures, *C*_1_ and *C*_2_, and *A* is the vesicle area. All our simulations are performed in constant *v* ensemble, where the fixed vesicle area, *A* = *A*_0_, identical for all vesicles, is preserved using a harmonic potential given by

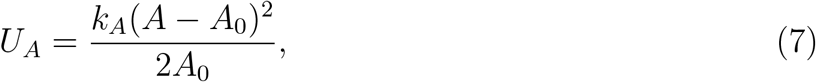

where *k*_*A*_ is the area conservation coefficient. To generate vesicles with given values of *v* for each simulation, the vesicle volume *V* is also constrained to remain nearly constant at target values *V*_0_ using the harmonic potential,

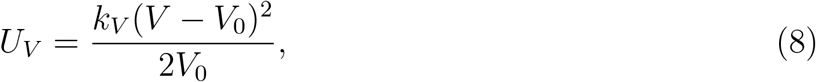

where *k*_*V*_ is the volume conservation coefficient. The links of the triangulated vesicles experience a bond potential which controls the link length. This potential, consists of a repulsive core (*U*_*r*_) and an attractive tail (*U*_*a*_), each taken to have a half-harmonic form given by:

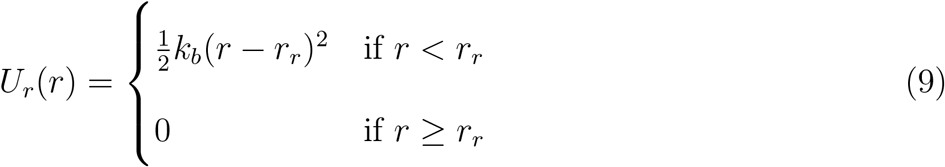

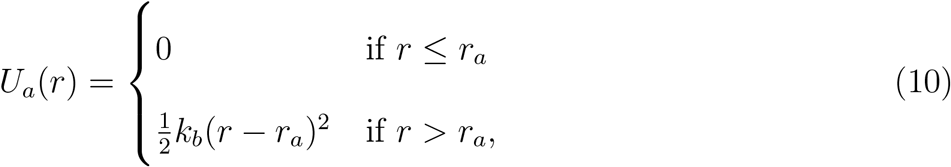

where *k*_*b*_ is the bond stiffness, and *r*_*a*_ and *r*_*r*_ are the potential cutoff lengths expressed in terms of *l*_0_ (see Table S1). Here, *l*_0_ denotes the side length of identical equilateral triangles whose total area equals the reference vesicle area *A*_0_. In our simulations with relatively low Pe ≤ 15, these half-harmonic potentials keep the lengths of the vesicle edges withing a range comparable to similar bond potentials.

### Membrane-membrane interactions

Small vesicle volumes typically lead to the formation of long, narrow tubes, thin sheets, and thin cup-like structures, where distant membrane segments may come into close proximity. In particular, opposing bilayers are found in close contact in sheet-like and cup-like morphologies. Additionally, different sections of a single long tube or two adjacent tubes in a branched structure may come into close proximity. While the latter case (tubular membranes) does not pose computational issues, intersecting apposing membranes lead to an underestimation of *v*. To prevent this, we apply a pairwise interaction potential, *U*_*vv*_(*r*) = *U*_*bv*_(*r*) (Eq. 4), between each pair of non-neighboring vertices of the vesicle. This computationally expensive interaction potential prevents membrane self-intersection, ensuring an accurate calculation of *v* and maintaining consistency with actual membranes, which do not exhibit self-intersection.

The sum of all these interaction potentials, along with *U*_*bv*_, which represents bead-vertex repulsive forces, and *U*_*vv*_, which prevents membrane self-intersection, gives rise to the total interaction potential, *U*_*vtot*_. The vesicle dynamics is modeled by the Langevin equation:

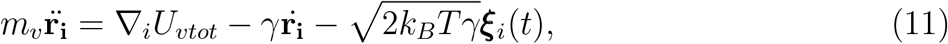

where *i* = 1..*N*_*v*_ runs over all vertices, *m*_*v*_ is the vertex mass, and other parameters are defined similar to Eq. 5 for filament beads. A Verlet algorithm was used to integrate equations of motion, Eqs. 5 and 11.

### Membrane fluidity

The triangulated vesicle model introduced so far represents a vesicle as a three dimensional network with a fixed connectivity between the network nodes as vesicle vertices. The network connectivity does not allow the vertices to freely move inside the vesicle plane. Actual membranes are however fluid in the sense that lipid molecules can easily diffuse inside the membrane plane. To fulfill membrane fluidity the network connectivity needs to be broken regularly in order to allow vertices freely diffuse in the vesicle membrane. This is done by flipping the vesicle bonds with a time frequency *ω* which replaces the two triangles sharing the edge by two new triangles. During flipping all vesicle bonds are flipped by a probability *q* where the flipping acceptance is decided by a Monte Carlo algorithm. The total change in the vesicle energy is calculated as the sum Δ*U* = Δ*U*_*bend*_ + Δ*U*_*A*_ + Δ*U*_*V*_ + Δ*U*_*r*_ + Δ*U*_*a*_ upon bond flipping. Bond flipping is then always accepted if it is energetically favorable with Δ*U <* 0 and otherwise with the probability of exp[−Δ*U/*(*k*_*B*_*T*)].

### Vesicle shape

To distinguish vesicle shapes, we use a vesicle shape index, *α* = Δ*a/*Δ*a*_*T*_, which is calculated from the membrane asymmetry Δ*a* of the vesicle and the membrane asymmetry Δ*a*_*T*_ of a cylindrical tube at the same volume. To estimate Δ*a* for sheet-like and tubular vesicles at each *v*, we use simplified axisymmetric cylindrical tube and toroidal sheet structures, as shown in Fig. S7A. The cylindrical tube is modeled as a straight cylinder of radius *r*_*T*_ and length *L*_*T*_ = *br*_*T*_, capped by two spherical caps at both ends (Fig. S7A right). Its surface area and volume are given by:

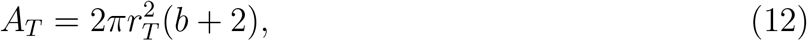

and

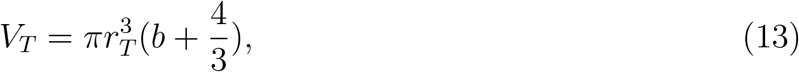

leading to a reduced volume,

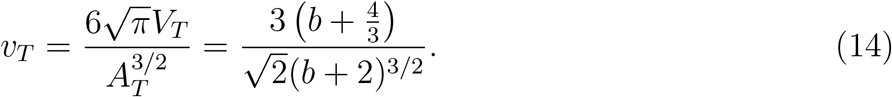

The membrane asymmetry of the tube is obtained from:

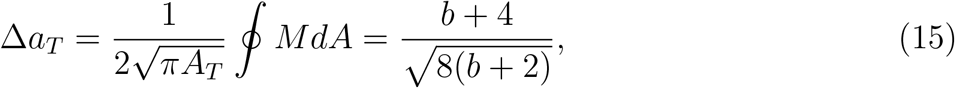

where we use *M* = 1*/r*_*T*_ (*C*_1_ = *C*_2_ = 1*/r*_*T*_) for the spherical caps and *M* = 1*/*(2*r*_*T*_) (*C*_1_ = 0, *C*_2_ = 1*/r*_*T*_) for the cylindrical section. We then use Eqs. 14 and 15 to express Δ*a*_*T*_ as a function of *v*, which in turn allows us to determine *α* for any vesicle shape at a given *v*. Table 1 presents the different values of *v* used in our simulations, along with the corresponding values of Δ*a*_*S*_ and Δ*a*_*T*_, given by the left and right curves in Fig. S7B for sheet-like and tubular vesicles, respectively. The left curve in Fig. S7B, which corresponds to toroidal sheets, is shown as white dashed lines in Figs. 1B, 4A, 5, S2, and S5. To demonstrate that *α* obtained in this way serves as an accurate measure for defining vesicle shape, we introduce a simplified axisymmetric sheet vesicle model. This shape consists of a toroidal surface with radius of curvature *r*_2_, enclosed by two circular planes of radius *r*_1_ = *cr*_2_ on both sides (Fig. S7A left). The area and volume of this sheet are given by:

**Table 1:** Membrane asymmetry of sheets and tubes versus reduced volume.

| $v$ | 0.25 | 0.3 | 0.35 | 0.4 | 0.45 | 0.5 | 0.6 | 0.7 | 0.8 |
| --- | --- | --- | --- | --- | --- | --- | --- | --- | --- |
| $\Delta a_S$ | 1.083 | 1.078 | 1.072 | 1.067 | 1.061 | 1.056 | 1.044 | 1.033 | 1.022 |
| $\Delta a_T$ | 3.06 | 2.57 | 2.22 | 1.96 | 1.77 | 1.61 | 1.39 | 1.23 | 1.13 |

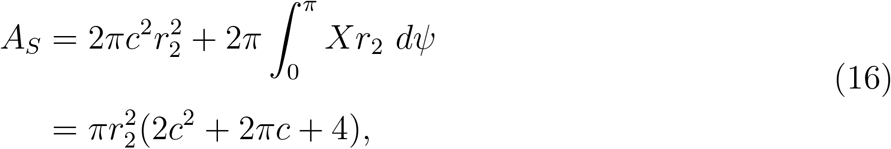

and

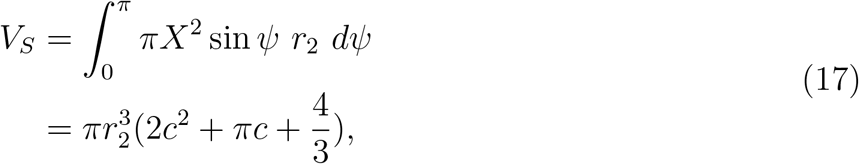

where *ψ* is the angle between the tangent line to the planar sheet contour and the horizontal axis and *X* = *r*_1_ + *r*_2_ sin *ψ* is the radial distance of a point on the contour form the axis of symmetry,^35,36,62^ Fig. S7A left. This leads to a reduced volume,

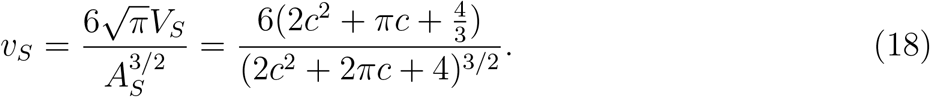

The membrane asymmetry of the sheet is obtained as:

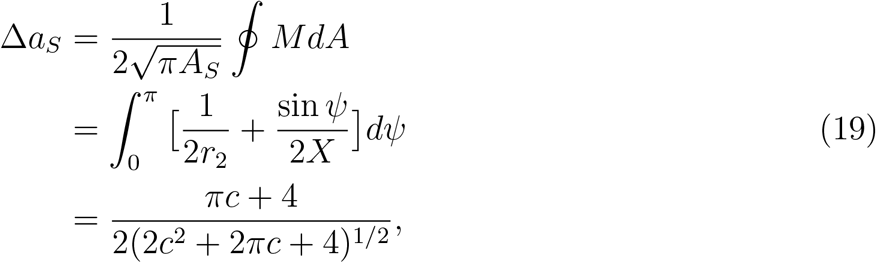

where *dA* = 2*πXds* = 2*πXrdψ* is expressed in terms of the arc length *s* on planar membrane contour as shown in Fig. S7A left. Equations 18 and 19 are used to compute Δ*a*_*S*_ at each *v*, which is then used to determine *α* for the sheet vesicle, as indicated by the left curve in Fig. S7B and the white dashed lines in Figs. 1B, 4A, 5, S2, and S5.

### Membrane tension

We calculate membrane tension using the virial expansion.^51,63^ The virial contribution from the global area and volume constraints is given by

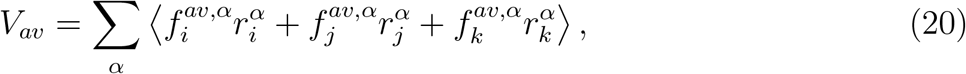

where 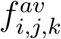 are the constraint forces acting on the vertices *i, j*, and *k* of a triangular membrane element, and *α* = *x, y, z* denotes the Cartesian coordinates. The virial contribution from the elastic bond forces *f*^*b*^ acting on the vesicle edges (links) is

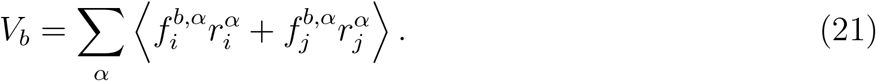

The total virial contribution from all forces on each vertex is then given by *V* = *V*_*av*_*/*3+ *V*_*b*_*/*2. Membrane tension is calculated via spatial and temporal averaging of the local stresses:

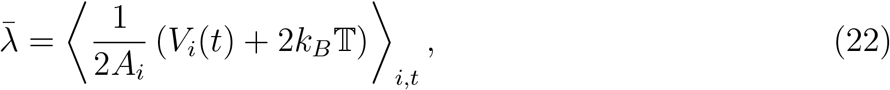

where *V*_*i*_(*t*) is the total virial contribution at vertex *i* at time *t*, and *A*_*i*_ is the area assigned to vertex *i*, approximately one-third of the total area of the surrounding triangles. The factor of two accounts for the two-dimensional nature of the membrane. The kinetic contribution 2*k*_*B*_*T* is estimated from the equipartition theorem. Contributions from the bending energy are neglected, as the associated forces act perpendicular to the membrane surface and do not contribute to in-plane tension.

## Conclusion

We explore dynamic shape remodeling of vesicles driven by the combined local effects of internally confined active filaments and the global effects of osmotic pressure, which regulates large-scale macroscopic shape transformations. In addition to the three fundamental vesicle shapes—tubes, sheets, and cups—which also occur in equilibrium in the absence of active elements, we observe intrinsically unstable hybrid structures such as branched tubes, sheet-tubes, pearled tubes, cup-tubes, and other compartmentalized morphologies like pear vesicles and sphere-tubes. These unstable hybrid vesicles are characterized by continuous dynamic restructuring, in which their constituent shape components interconvert while maintaining nearly constant proportions. For example, branched tubular networks reorganize by varying the number of tubular junctions while keeping the total tubular length nearly constant. Similarly, sheet-tubes restructure by changing the number of sheet-like and tubular segments while preserving the relative proportion of each component.

These dynamic structures are distinct from the non-equilibrium stationary shapes reported for similar vesicles containing confined active filaments in the absence of osmotic stress.^48^ They also differ from the stationary vesicle shapes—typically tubes, sheets, and cups—observed under internal active Brownian particles and osmotic stress, which transform into one another as particle mobility varies.^43^ For instance, the branched tubes we observe do not resemble the membrane tethers reported for vesicles containing both normal and sticky active particles. ^16,50,51^ These tethers typically form at relatively high values of both Pe and *v*, and are characterized by their very small diameter, comparable to the size of a single Brownian particle. In contrast, the branched tubular structures reported here exhibit a nearly uniform tube diameter that increases with *v*. Moreover, whereas membrane tethers are primarily observed at Pe ≲ 800, our branched tubes emerge at much lower values of Pe ≤ 15.^48^ Notably, branched tubes are more abundant at low Pe and are limited to small volumes *v* ≤ 0.4. While increasing Pe promotes tether formation, the occurrence of branched tubes increases with decreasing Pe.

Remarkably, the dynamic reorganization of these unstable hybrid structures accelerates as Pe decreases. Moreover, at relatively small Pe, the vesicles exhibit low membrane tensions that remain nearly constant across different values of Pe. This suggests that the dynamic restructuring of these floppy vesicles is governed more by osmotic effects than by filament mobility. Overall, the regime of low filament mobility inside floppy vesicles gives rise to unstable hybrid morphologies—such as branched tubes, sheet-tubes, and pearled vesicles—that continuously reorganize their shapes at rates that increase as Pe decreases.

We demonstrate that both the formation and dynamics of these unstable hybrid structures strongly depend on the anisotropic nature of the active filaments. Reducing filament stiffness at fixed length, or decreasing filament length (i.e., number of beads) at fixed stiffness, results in less anisotropic filaments with smaller P. Dynamically reorganizing hybrid vesicles—such as branched tubes and sheet-tubes—become less abundant as filament anisotropy decreases, and eventually vanish when filaments become fully crumpled or reduce to singlebead isotropic active particles (spherical particles), giving rise to cup-like vesicles. Our results thus highlight the importance of the filamentous nature of the cytoskeletal network in forming non-equilibrium dynamic structures of the cellular membrane. In particular, the dynamic branched tubes and sheet-tubes resemble cellular protrusions such as filopodia and lamellipodia, as well as the membrane morphologies observed in organelles like the endoplasmic reticulum and mitochondria.

## Data availability

All relevant data supporting the findings of this study are included in the article and its Supplementary Information.

## Acknowledgement

AHB acknowledges support from the Max Planck Society within the framework of Max Planck Partner Group, and from the European Molecular Biology Organization, grant EMBO IG 5032.

## Author contribution

AHB designed research; AKS and AHB performed simulations; AKS and AHB analyzed the data. AHB wrote the manuscript with assistance from AKS.

## Supporting Information Available

The Supplementary information includes supplementary figures, tables, and videos.

## Supplementary Information

This Supplementary Information contains:

### 1. Supplementary Figures

**Fig. S1.**
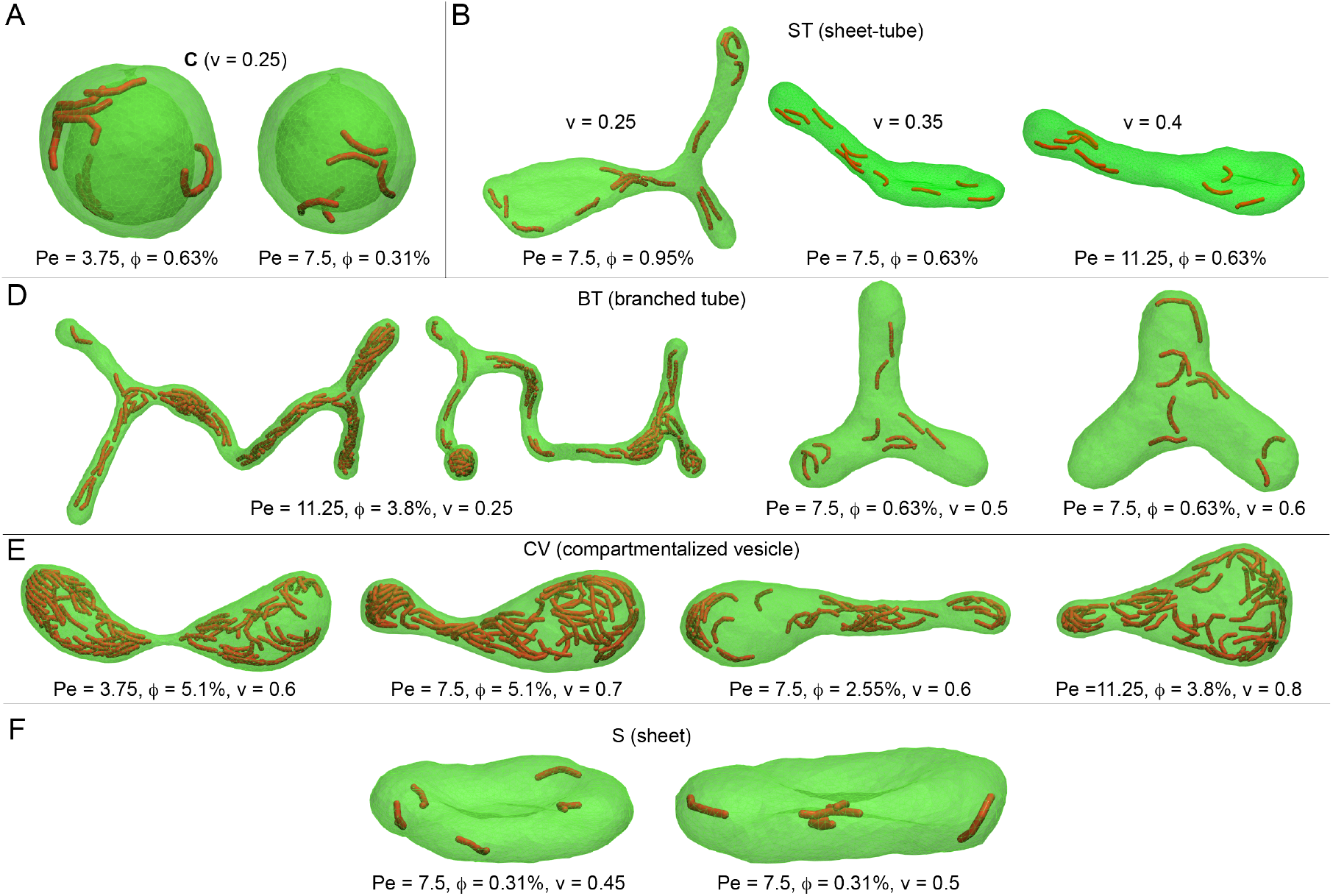
Vesicle morphologies formed by internal active filaments. Vesicle morphologies formed by internal active filaments. Different vesicle shapes are shown with their corresponding simulations times from left to right in each panel. (A) Cups (40 s and 50 s). (B) Sheet-tubes (50 s for all three snapshots). (D) Branched tubes at low and high volumes (36.8 s, 41.7 s, 50 s, and 20.4 s). (E) Compartmentalized vesicles including pearled tube (left) and pear-shaped vesicles (right) (33.6 s, 50 s, 50 s, and 50 s). (F) Sheets (50 s for both snapshots).

**Fig. S2.**
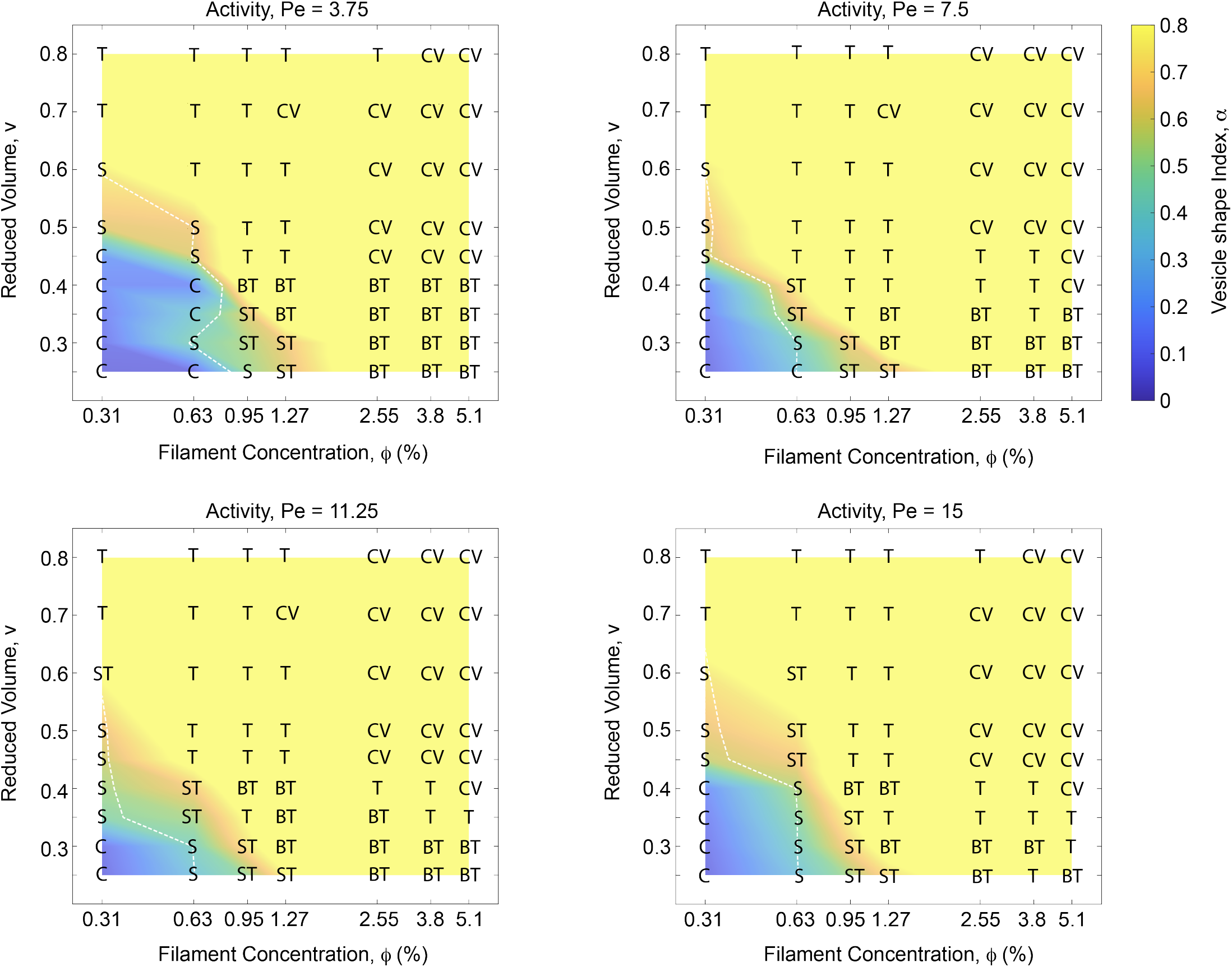
Morphology diagram of active vesicles Morphology diagram of active vesicles. Vesicle shapes with varying *ϕ, v*, and Pe for relatively stiff filaments (χ = 25) of intermediate length (ℒ = 6.12). Vesicle shapes are characterized by the shape index *α*, which is based on membrane asymmetry (Δ*a*) and is color-mapped onto the shape diagram. The white dashed line indicates toroidal sheets predicted by the theoretical vesicle model at different values of *v* (see Methods).

**Fig. S3.**
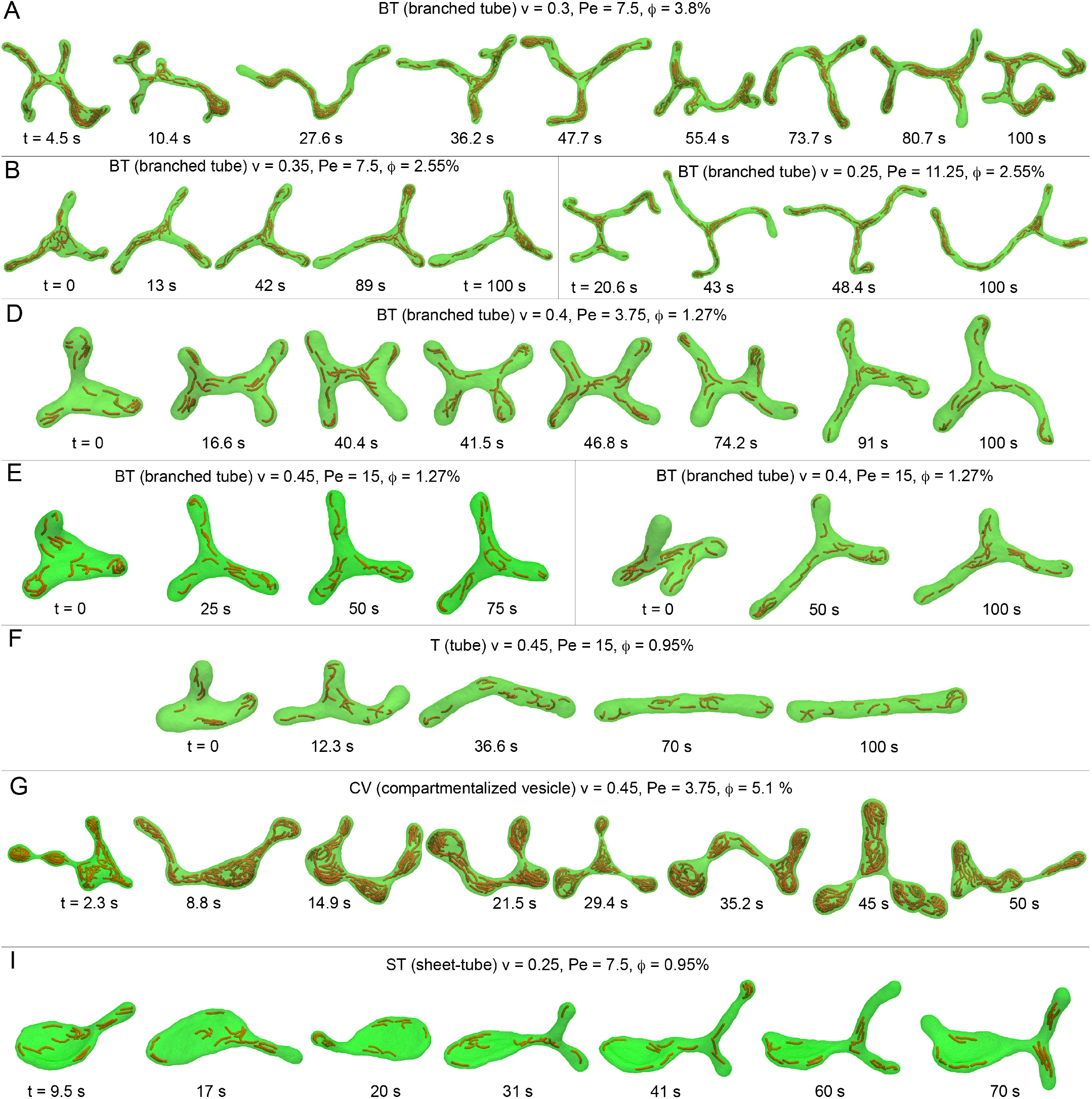
Temporal evolution of dynamically reorganizing active vesicles Temporal evolution of dynamically reorganizing active vesicles. (A-E) Branched tubular networks with different values of *v*, continuously reorganize their structures by extending and retracting tubes, forming a variable number of three-way tubular junctions. (F) Unstable short-lived branched tube at relatively large *v* = 0.45 transforming to a stable tube. (G) Temporal evolution of a highly dynamic compartmentalized vesicle, composed of restructuring compartments, at a relatively large *ϕ* over 50 seconds. (I) A sheet-tube (ST) dynamically reorganizes its shape by transitioning from one to two tubular segments over 60 seconds.

**Fig. S4.**
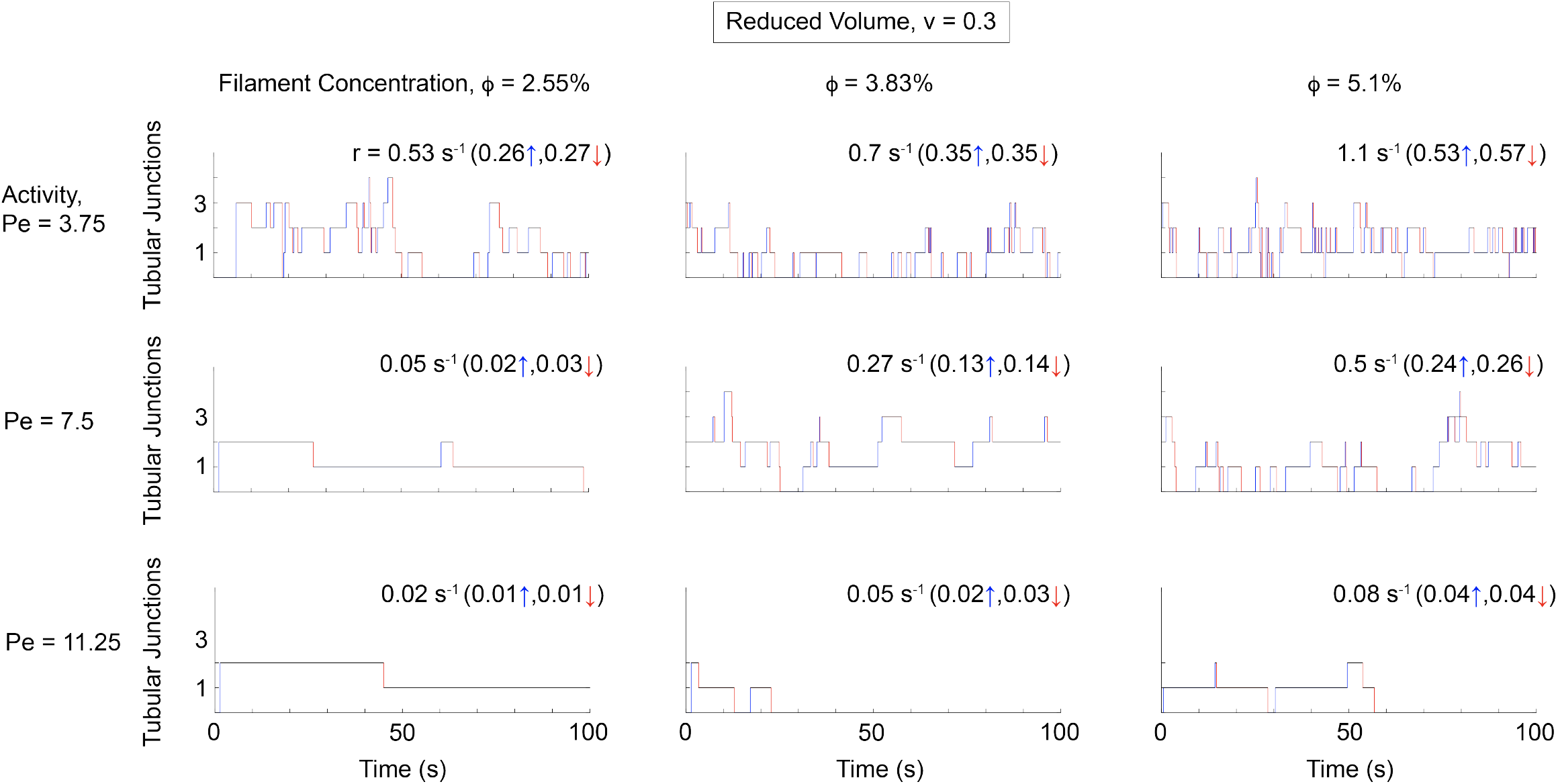
Dynamic reorganization of highly branched tubular networks Dynamic reorganization of highly branched tubular networks. Temporal evolution of the number of three-way tubular junctions for different values of Pe and *ϕ* at *v* = 0.3. The total rate of change in the number of tubular junctions is given by the sum of the tube formation rate (blue lines) and the tube retraction rate (red lines). The rate of change in tubular junctions increases with *ϕ* for a fixed Pe, while it decreases with Pe at a constant *ϕ*. All branched tubes exhibit nearly identical formation and retraction rates, implying that tube formation and retraction occur with almost equal frequency.

**Fig. S5.**
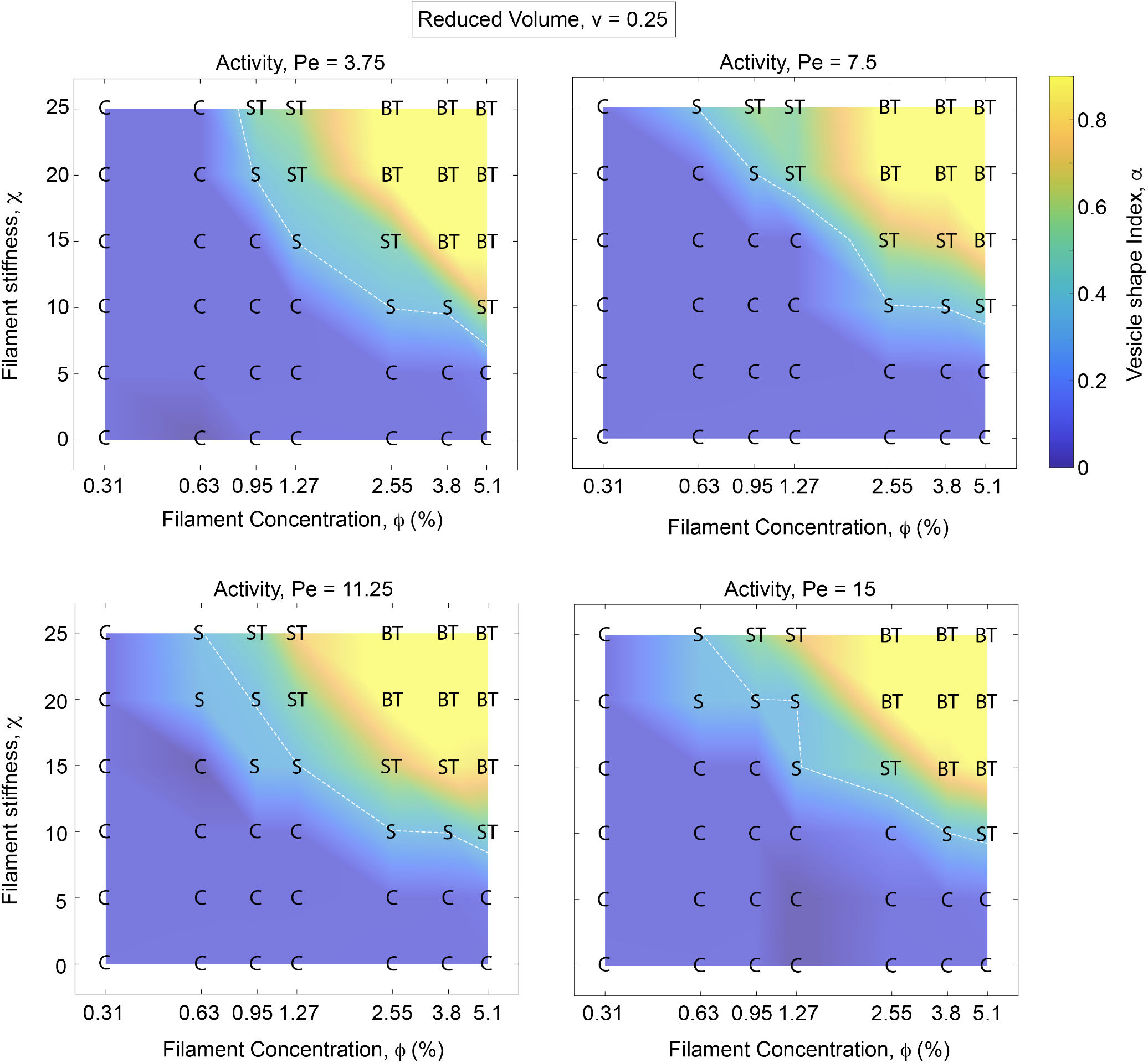
Morphological behavior of vesicles with active filaments at varying *χ* Morphological behavior of vesicles with active filaments at varying χ. Vesicle shapes are obtained for *v* = 0.25 and different values of Pe. Lower filament stiffness delays the transition from cups to sheets, sheet-tube structures, and eventually to branched tubes at higher values of *ϕ*. Reduced stiffness also leads to the formation of crumpled filaments, transforming branched tubes into cups for χ ≤ 5. The white dashed line indicates toroidal sheets predicted by the theoretical vesicle model at different values of *v* (see Methods). Vesicle morphologies show only weak sensitivity to Pe.

**Fig. S6.**
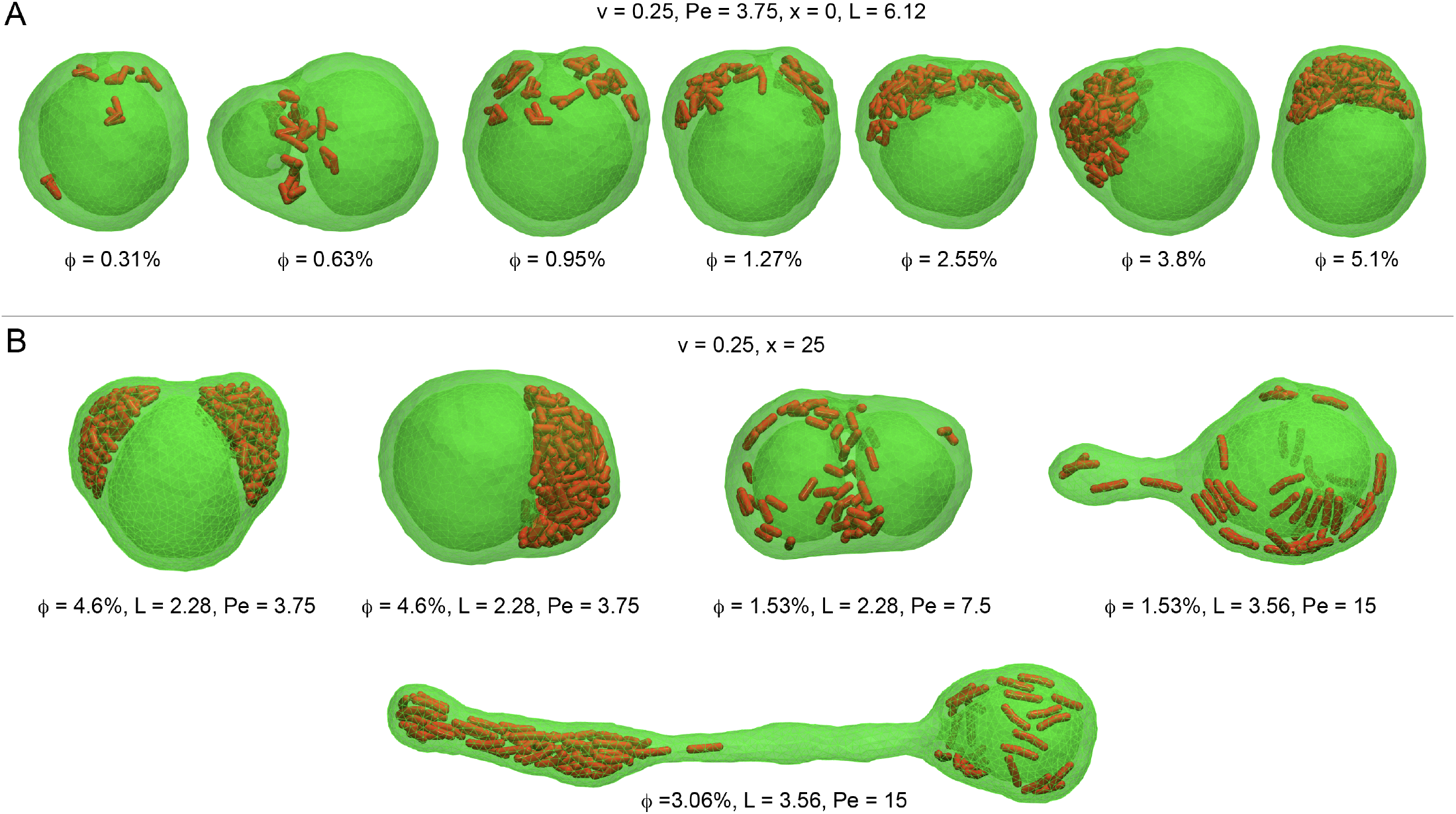
Vesicle morphologies for varying *L* and *χ* Vesicle morphologies for varying ℒ and χ. (A) A variety of cup-like vesicles formed by active filaments with negligible bending stiffness (χ), shown for different values of *ϕ* at fixed *v* = 0.25. (B) Distinct cup-like variants, including compartmentalized cups, double-cups, and cup-tubes (comprising both cup-like and tubular segments), observed for stiff active filaments with varying filament lengths ℒ, ranging from ℒ = 1 (corresponding to spherical active beads) to ℒ *>* 1 (extended active filaments). All snapshots correspond to *t* = 50 s.

**Fig. S7.**
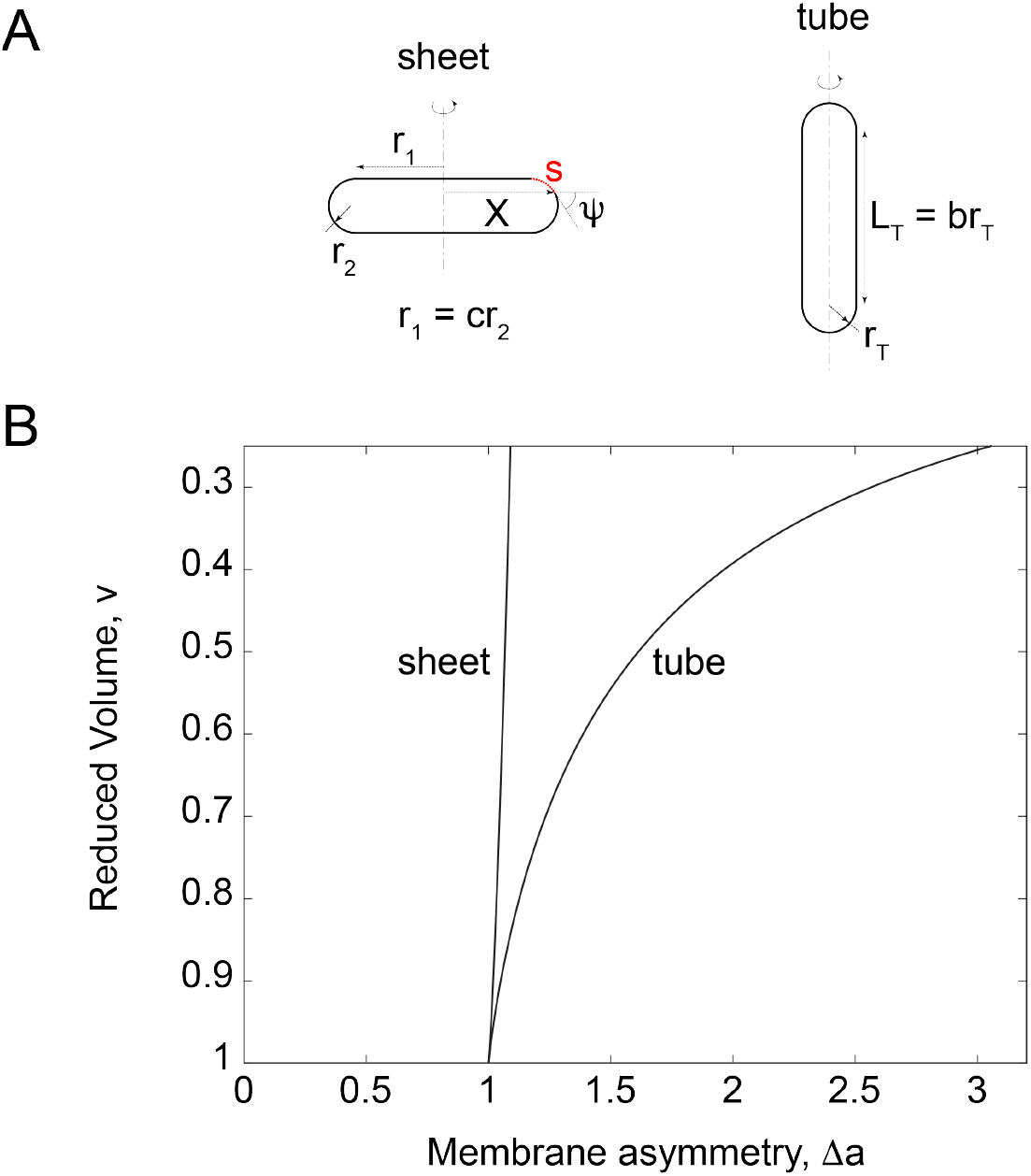
Theoretical vesicle models Theoretical vesicle models. (A) Schematics of axisymmetric vesicle shapes: a toroidal sheet (left) and cylindrical tubes (right). (B) Reduced volume of toroidal sheets and cylindrical tubes plotted as a function of membrane asymmetry, Δ*a*. A straight line is fitted to the toroidal sheet data, defining the white dashed lines in Figs. 1, 4, 5, S2, and S5. The vesicle shape index, *α* = Δ*a/*Δ*a*_*T*_, is calculated using the membrane asymmetry of the cylindrical tube, Δ*a*_*T*_ (right curve), at each value of *v*.

**Fig. S8.**
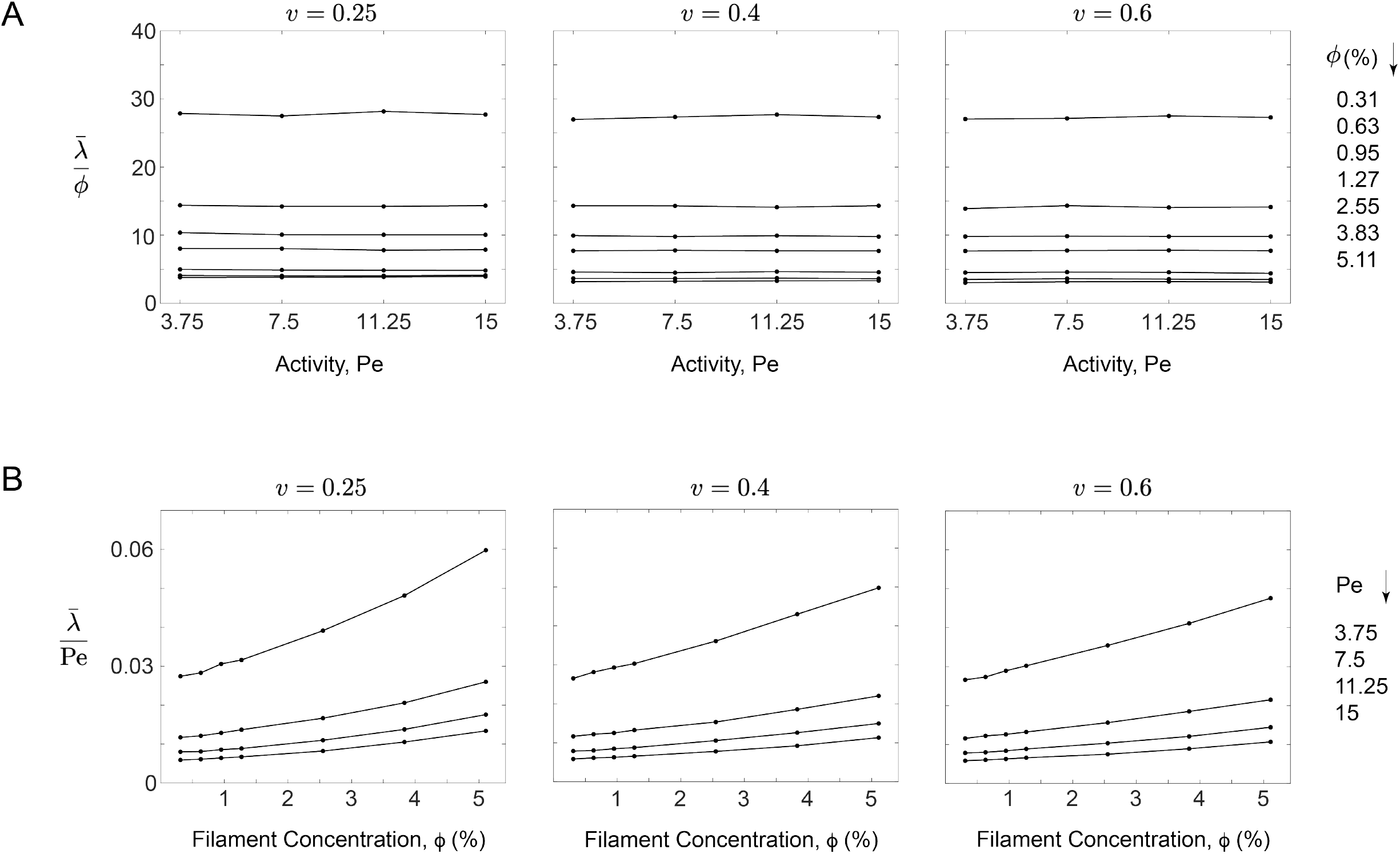
Membrane tension in active vesicles Membrane tension in active vesicles. (A) Average membrane tension as a function of Pe for different values of *ϕ*. (B) Average membrane tension as a function of *ϕ* for different values of Pe.

### 2. Supplementary Table

**Tab. S1:** Principal parameters

### 3. Legends of Supplementary Videos

**Movie S1:** Dynamic shape remodeling of a branched tube at low volume

**Movie S2:** Dynamic shape remodeling of a branched tube at intermediate volume

**Movie S3:** Dynamic shape remodeling of a sheet-tube

**Table S1:** Principal parameters.

| Parameters | Model units | Physical units |
| --- | --- | --- |
| Basic properties |  |  |
| reference vesicle radius $R$ | 0.62035 | 1 $\mu\text{m}$ |
| time scale $\tau = \gamma R^2 / \kappa$ | $1.925 \times 10^5$ | 0.1925 s |
| thermal energy unit $k_B T$ | 1 | $4.14 \times 10^{-21}$ J |
| System parameters |  |  |
| Péclet number $\text{Pe} = \sigma f_p / k_B T$ | 3.75 - 15 | 3.75 - 15 |
| number of beads per filament $N_{bf}$ | 1 - 10 | 1 - 10 |
| number of beads $N_b = N_f \times N_{bf}$ | 25 - 400 | 25 - 400 |
| filament concentration $\phi = N_f N_{bf} (\sigma / 2R)^3$ | 0.31 - 6.4 | 0.31 - 6.4 |
| desired vesicle volume $V_0$ | 0.25 - 0.8 | 1.05 - 3.35 $\mu\text{m}^3$ |
| filament length $\mathcal{L} = (N_b - 1)r_0 + \sigma$ | 1 - 12.52 | 1 - 12.52 |
| filament stiffness $\chi = \kappa_f / \kappa$ | 0 - 25 | 0 - 25 |
| Vesicle properties |  |  |
| number of vertices $N_v$ | 2562 | 2562 |
| bending rigidity $\kappa$ | $20k_B T$ | $8.28 \times 10^{-20}$ J |
| equilibrium bond length $l_0$ | $4R \sqrt{\frac{\pi}{N_t \sqrt{3}}}$ | 0.075 $\mu\text{m}$ |
| potential cutoff $r_r$ | $0.95l_0$ | 0.0715 $\mu\text{m}$ |
| potential cutoff $r_a$ | $1.05l_0$ | 0.079 $\mu\text{m}$ |
| bond stiffness $k_b$ | $11.54k_B T / R^2$ | $4.78 \times 10^{-17}$ J.$\mu\text{m}^{-2}$ |
| desired vesicle area $A_0$ | $4\pi R^2$ | 12.57 $\mu\text{m}^2$ |
| area conservation coefficient $k_A$ | $3.72 \times 10^5 k_B T / R^2$ | $1.54 \times 10^{-15}$ J.$\mu\text{m}^{-2}$ |
| volume conservation coefficient $k_V$ | $2.38 \times 10^6 k_B T / R^3$ | $9.88 \times 10^{-15}$ J.$\mu\text{m}^{-3}$ |
| friction coefficient $\gamma$ | $20k_B T \tau / R^2$ | $1.59 \times 10^{-20}$ J.s$\mu\text{m}^{-2}$ |
| Active filament properties |  |  |
| effective bead diameter $\sigma$ | $R/10$ | 0.1 $\mu\text{m}$ |
| equilibrium filament bond length $r_0$ | $1.28\sigma$ | 0.129 $\mu\text{m}$ |
| potential coefficient $k_{bb}$ | $1.15 \times 10^6 k_B T / R^2$ | $4.78 \times 10^{-15}$ J.$\mu\text{m}^{-2}$ |
| filament bond stiffness $k_{fb}$ | $1.15 \times 10^4 k_B T / R^2$ | $4.78 \times 10^{-17}$ J.$\mu\text{m}^{-2}$ |

**Movie S1. Dynamic shape remodeling of a branched tube at low volume**. Dynamic shape remodeling of the branched tube at *v* = 0.25 over 100 seconds, driven by internal active filaments. The movie corresponds to the snapshots shown in Fig. 2A. At the smallest filament mobility (Pe = 3.75), this branched tube rapidly reorganizes its structure by dynamically varying the number of tubular junctions, as illustrated in the top-left panel of Fig. 3A.

**Movie S2. Dynamic shape remodeling of a branched tube at intermediate volume**. Dynamic shape remodeling of the branched tube at *v* = 0.35 over 100 seconds, as shown in the snapshots in Fig. 2B. Compared to the branched tube in Movie. S1, which has a smaller *v*, this tube—with the same Pe—exhibits wider diameters and slower dynamic reorganization.

**Movie S3. Dynamic shape remodeling of a sheet-tube**. Dynamic shape transitions of a sheet-tube at *v* = 0.25 over 100 seconds, corresponding to the snapshots in Fig. 2D. Despite continuous interconversion between sheet-like and tubular sections, the area proportion of each component remains nearly constant. Although the number of tubes and their individual lengths fluctuate, both the total tubular length and the tube diameter remain approximately unchanged.

